# Probiotic Associated Therapeutic Curli Hybrids (PATCH)

**DOI:** 10.1101/464966

**Authors:** Pichet Praveschotinunt, Anna M. Duraj-Thatte, Ilia Gelfat, Franziska Bahl, David B. Chou, Neel S. Joshi

## Abstract

There is an unmet need for new treatment methods for inflammatory bowel disease (IBD) that can reliably maintain remission without leading to detrimental side effects. Beneficial bacteria have been utilized as an alternative treatment for IBD albeit with low efficacy. We genetically engineered *Escherichia coli* Nissle 1917 (EcN) to create an anti-inflammatory fibrous matrix *in situ*. This matrix consists of EcN-produced curli nanofibers displaying trefoil factors (TFFs), known to promote intestinal barrier function and epithelial restitution. We confirmed that engineered EcN was able to secrete the curli-fused TFFs *in vitro* and *in vivo*, and was non-pathogenic. We observed an enhanced protective effect of engineered EcN against dextran sodium sulfate induced colitis in mice, associated with barrier function reinforcement and immunomodulation. This work sets the foundation for the development of a novel therapeutic platform in which the *in situ* production of a therapeutic protein matrix from beneficial bacteria can be exploited.

## Introduction

Inflammatory Bowel Disease (IBD) describes a group of autoimmune diseases that cause chronic inflammation of the small and large intestine. This group of diseases affects about 3 million adults in the U.S. (*1*). A complete picture of IBD etiology remains a subject of intense research and debate, and the development of the disease has been linked to multiple factors, including dysregulated immune responses, genetic predisposition, and an altered balance of microbiota (i.e. dysbiosis) (*2*). However, the complex interplay between these factors has greatly hindered the development of effective therapies. Conventional IBD treatment relies on pharmacological interventions that scale with the severity of the disease, starting with aminosalicylates and antibiotics, and proceeding to corticosteroids and immunosuppressants, with the goal of dampening inflammation by influencing host biology or by decreasing the chance of bacterial infection of mucosal wounds. The consequences of disease flare-ups can be severe and mount over time, leading to 23-45% of ulcerative colitis patients and 75% of Crohn’s disease patients requiring surgical removal of portions of their gastrointestinal (GI) tract at some point in their lives (*3*). Therefore, the ability to induce deep remission and sustain it indefinitely is the long-term goal of IBD treatments.

Some early successes with therapies that target tumor necrosis factor (TNF) initially indicated a promising future for biologics in the treatment of severe cases of IBD. However, progress in the development of new therapeutics has been slow, with several notable clinical failures arising from drug candidates with strong results in small animal models (*4*). A common theme in these failures is that the efficacy for a given treatment varies according to patient sub-population or environmental factors (e.g. diet, social behaviors). Given the heterogeneity in disease etiology, it is likely that multi-pronged and perhaps patient-specific management strategies will be necessary to achieve effective clinical outcomes (*5*).

An emerging area of IBD research deals with gut microbes. Although it remains unclear whether dysbiosis can incite disease, it is clear that global reductions in gut microbiome diversity are correlated with IBD severity (*6*). Furthermore, it is also clear that many microbiota can exacerbate IBD-associated inflammation via compromised epithelial barrier function (*7*). These factors have led to the exploration of living bacteria as therapeutic entities that can be delivered orally (i.e. “bugs as drugs”). Several naturally occurring commensal and beneficial strains have been explored as therapeutics, with limited success mostly stemming from their low potency and inability to persist in the GI tract (*8*, *9*). Genetically engineered microbes have also been explored, mostly as a means to secrete biologic drugs (e.g. IL-10, anti-TNF) locally in the colon (*10*-*12*). Many such efforts have also shown high efficacy in animal models but have yet to yield clinical successes, in part because of difficulties in achieving and maintaining sufficiently high concentrations of the therapeutic molecule at the site of disease. Indeed, concerns have been raised about the compatibility of this strategy with immunomodulatory biologics, since the mucosal epithelial barrier hinders their trafficking to their target cells in the lamina propria (*13*, *14*). Nevertheless, the promise of effective treatments that can be produced cheaply, delivered orally, and minimize systemic side effects has continued to fuel interest in microbes as therapeutics.

Here we present an alternative approach to engineered microbial therapies for IBD treatment. Instead of secreting soluble therapeutic proteins, we programmed bacteria to assemble a multivalent material decorated with anti-inflammatory domains in the gut. The displayed domains are designed to target the material to the mucosal layer of the epithelium and promote host processes that reinforce epithelial barrier function (Figure 1). The bacterially-produced scaffold for the living material is based on curli fibers, a common proteinaceous component of bacterial extracellular matrices. Hence, we refer to our approach as Probiotic Associated Therapeutic Curli Hybrids (PATCH). We demonstrate that PATCH is capable of ameliorating inflammation caused by dextran sodium sulfate (DSS) induced colitis in a mouse model.

**Figure 1.**
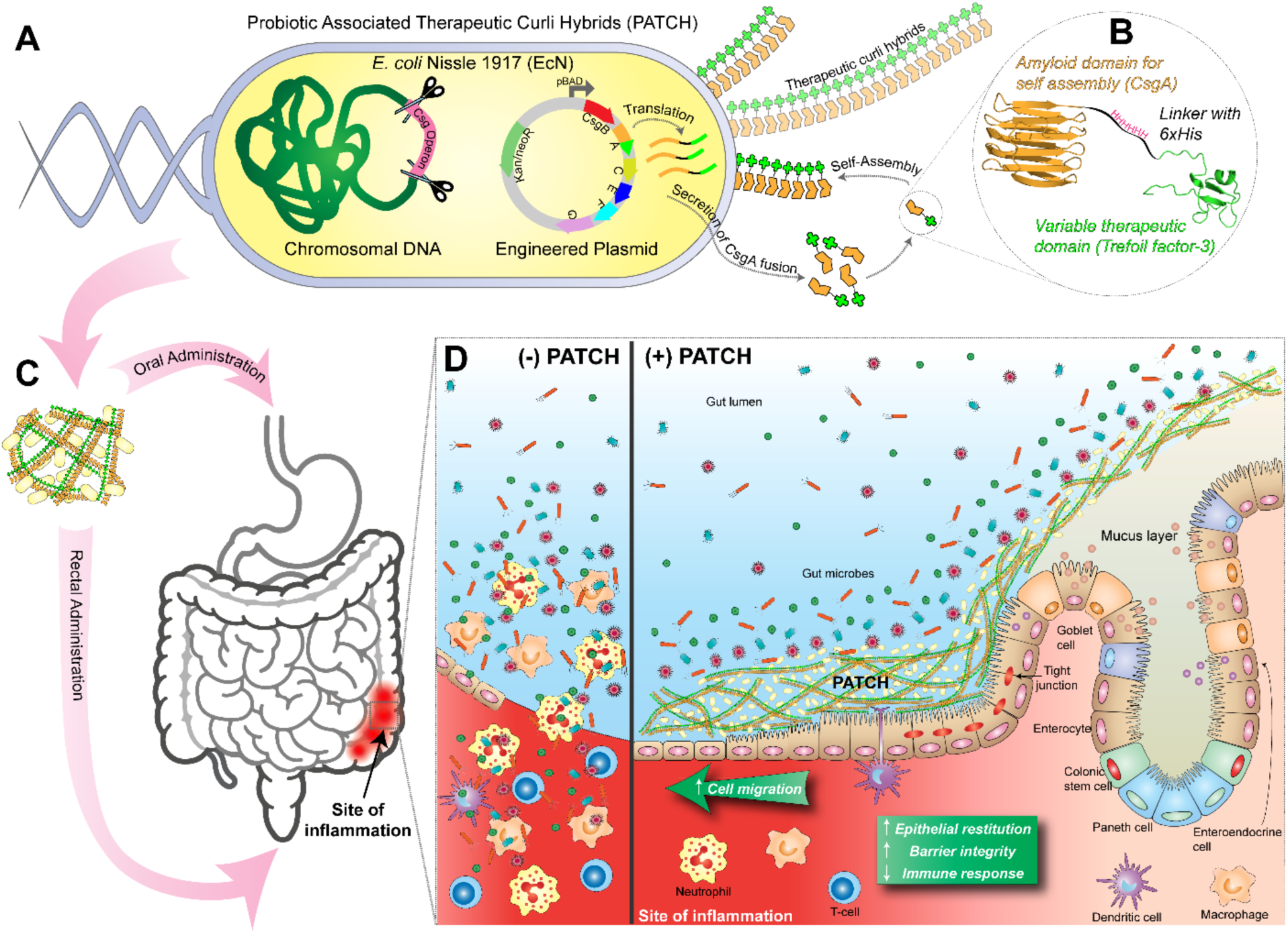
Probiotic Associated Therapeutic Curli Hybrids (PATCH). (A) Schematic overview of engineered curli production. Genetically engineered E. coli Nissle 1917 (EcN) with csg (curli) operon deletion (PBP8 strain) containing plasmids encoding a synthetic curli operon capable of producing chimeric CsgA proteins (yellow chevrons with appended bright green domains), which are secreted and self-assembled extracellularly into therapeutic curli hybrid fibers. (B) CsgA (yellow), the main proteinaceous component of E. coli biofilm matrix, was genetically fused to a therapeutic domain – in this case, TFF3 (PDB ID: 19ET, bright green), which is a cytokine secreted by mucus producing cells. The flexible linker (black) includes a 6xHis tag for detection purposes. (C) Engineered bacteria are produced in bulk before delivery to the subject via oral or rectal routes. A site of colonic inflammation is highlighted in red. (D) Interaction of PATCH and the colonic mucosa. Inflammatory lesions in IBD result in loss of colonic crypt structure, damage to epithelial tissue and compromised barrier integrity (left panel, (−) PATCH). The resulting invasion of luminal contents and recruitment of immune cells to the site exacerbates the local inflammation. The application of PATCH (right panel, (+) PATCH) reinforces barrier function, promotes epithelial restitution, and dampens inflammatory signaling to ameliorate IBD activity.

## Results

We used *E. coli* Nissle 1917 (EcN) as our cellular chassis for PATCH. EcN is well-studied, has a long track record of safety in humans, and is a popular starting point for engineered therapeutic microbe efforts because of its compatibility with canonical genetic engineering techniques for bacteria (*15*). In addition to its use as an over-the-counter supplement for general GI disorders, EcN has also been evaluated in comparison to mesalazine for maintaining remission in ulcerative colitis in randomized control trials (*16*). While EcN lead to some favorable outcomes, overall efficacy was low and relapse rates were high, impeding its use as a first-line treatment for IBD (*16*, *17*). Like other Enterbacteriaceae, EcN resides mostly in the colon, where it colocalizes with areas affected by many types of IBD (*18*). Moreover, facultative anaerobes like EcN are known to thrive in the highly oxidative environment of the inflamed GI tract (*19*), making EcN an ideal starting point for our engineering efforts.

We chose the trefoil factor (TFF) family of human cytokines as our bioactive domain for display on curli fibers. TFFs are small, 7-12 kDa proteins secreted by mucus producing cells in the GI tract and other mucosal surfaces, primarily to promote epithelial restitution (*20*). TFFs also reportedly have tumor suppressing, apoptosis blockading, and barrier function augmenting bioactivity, though the precise mechanisms for these effects are still not well understood (*20*, *21*). TFFs have been explored for IBD treatment, but oral delivery did not yield therapeutic outcomes, as they were found to adhere too strongly to the mucus layer of the small intestine (*11*). We sought to overcome this by tethering them to the curli fiber matrix after local production in the ileum, cecum, and colon.

### TFF-fused curli fibers are successfully secreted and assembled by engineered E. coli Nissle 1917

In order to implement the PATCH system, we created plasmid-based genetic constructs encoding for the self-assembling monomer unit of curli fibers (CsgA) fused to each of the three TFFs (TFF1-3). The TFFs were appended to the C-terminus of CsgA via a flexible glycine-serine linker containing and internal 6xHIS tag in a manner that we have previously shown to not interfere with extracellular secretion and self-assembly (*22*). The library of plasmids was designed such that each gene encoding a CsgA-TFF fusion was co-transcribed with the other genes necessary for effective curli secretion and assembly (*csgB*, *csgC*, *csgE*, *csgF*, *csgG*). Together, these formed a “synthetic curli operon” that was placed under the control of an inducible promoter (P_BAD_) in a pBbB8k plasmid backbone bearing a kanamycin selection marker (Figure 1). The inclusion of the other genes of the curli operon was necessary to increase secretion efficiency, because the curli genes in the EcN chromosome are downregulated at physiological temperature and osmolarity (*23*). Nevertheless, we also employed an EcN mutant in which all of the chromosomal curli genes were deleted (EcN Δ*csgBACDEFG*::*Cm^R^*, a.k.a PBP8) in order to preclude the possibility of curli fiber expression from native genes confounding our experimental results (*24*).

In order to confirm that curli fibers decorated with TFFs could be produced by EcN, as they can in laboratory strains of *E. coli* (*22*), we transformed EcN with the panel of synthetic curli plasmid constructs. The transformed cells were cultured at 37°C in high osmolarity media to mimic physiological conditions and induced with L-(+)-arabinose. A quantitative Congo Red binding (CR) assay, normally used for curli fiber detection (*22*, *25*), indicated that wild-type CsgA and all three CsgA-TFF fusions could be expressed and assembled into curli fibers under physiological conditions (Figure 2A). Extracellular assembly was further confirmed using whole-cell ELISA assays probing for the 6xHIS tag (Figure 2B). Scanning electron microscopy of the samples confirmed that recombinant wild-type CsgA and all the CsgA-TFF fusions assembled into nanofibrous structures resembling native curli fiber in appearance (Figure 2C-G). Curli production experiments performed with PBP8 led to similar trends. In some cases, basal expression of the *csgA* genes was observed without induction (Figure S1A, B). We have also confirmed the presence of the displayed TFF3 using similar whole cell ELISA assay (Figure 2H).

**Figure 2.**
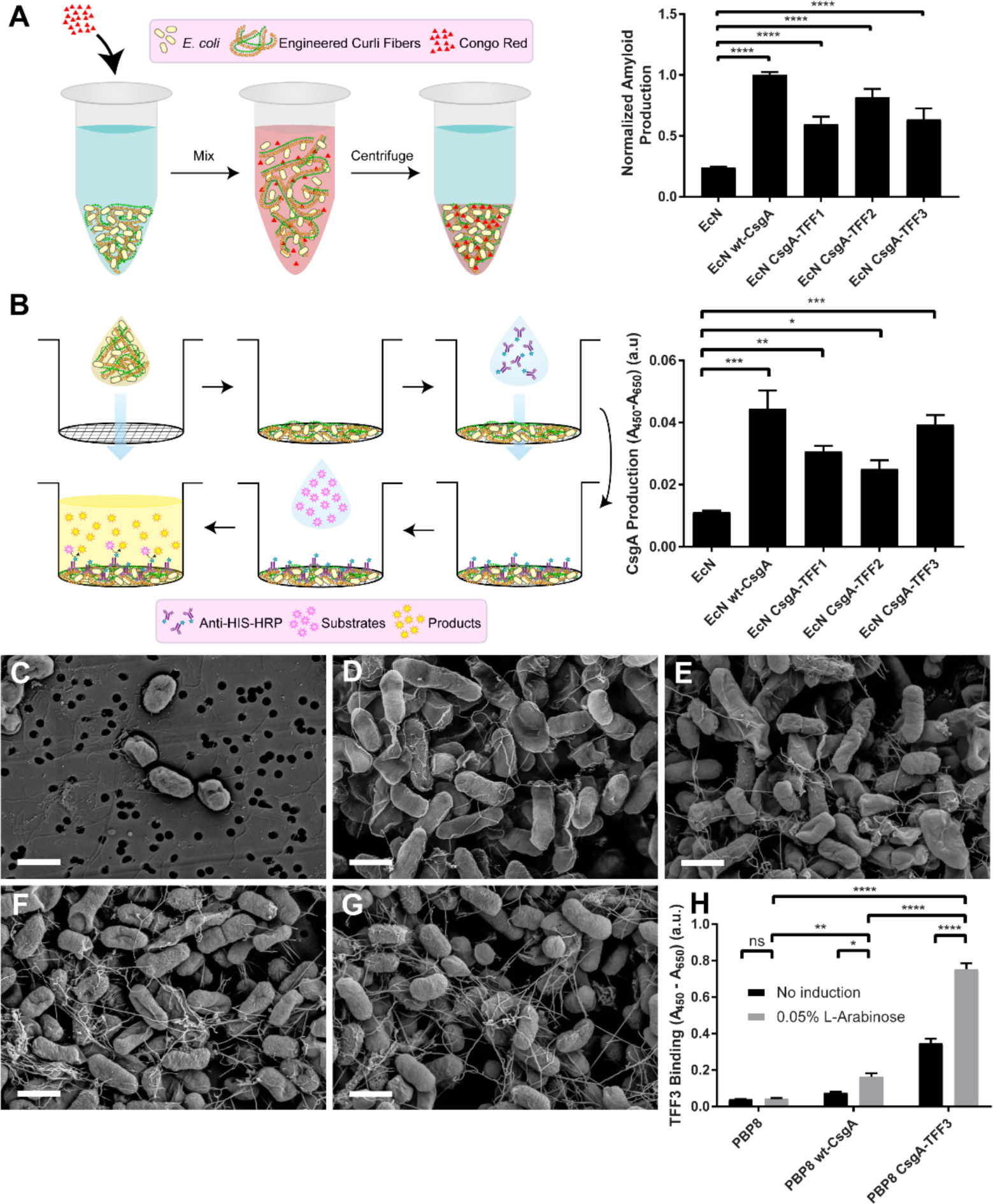
Production of curli fiber variants from engineered EcN. (A) Schematic of quantitative Congo Red (CR) binding assay (left). Normalized amyloid production of each EcN variant, as measured by CR binding assay (right). (B) Schematic of whole-cell filtration ELISA for the monitoring of modified curli production from bacterial culture (left). Relative CsgA production between EcN variants, derived from anti-6xHis antibody-based detection. (C-G) Scanning electron micrographs of EcN transformed with plasmids encoding various proteins: (C) GFP (D) wt-CsgA (E) CsgA-TFF1 (F) CsgA-TFF2 (G) CsgA-TFF3 (scale bar = 1 μm). (H) Relative TFF3 production (using anti-TFF3 antibody) of induced and non-induced PBP8 library. Data are represented as mean ± SEM. See materials and methods for statistics.

### Engineered curli fibers do not confer pathogenicity to EcN in vitro

We had previously demonstrated that the CsgA-TFF3 fusion, when produced by a laboratory strain of *E. coli*, could bind to mucins and promote mammalian cell migration in an *in vitro* model with a human colorectal adenocarcinoma cell line (Caco-2) (*22*). Before proceeding to *in vivo* studies, we wanted to confirm that modified curli fiber overproduction did not induce a pathogenic phenotype in PBP8. Therefore, we performed invasion and barrier function assays on Caco-2 cells grown to confluency in transwells. None of the EcN derived strains (i.e. PBPB8 – curli genes deleted, PBP8 wt-CsgA – expressing the wild-type CsgA sequence, PBP8 CsgA-TFF3 – expressing the CsgA-TFF3 fusion) exhibited increased invasiveness into polarized Caco-2 monolayers when compared to unmodified EcN (Figure 3). Invasion for Caco-2 monolayers was low across all groups in comparison to a positive control, *Salmonella typhimurium* (SL 1344) (Figure 3A). Similarly, in a translocation assay in which bacteria were collected from the basolateral chamber of the transwell (*26*), we observed essentially no translocation of any of the EcN derived strains (Figure 3B). We also monitored barrier function in the transwells as a function of bacterial strain. Trans-epithelial electrical resistance (TEER) measurements showed lower reductions TEER values for all of the EcN derived strains compared to *S. typhimurium* – 40-50% vs. 70%, respectively (Figure S2). We also tested barrier function *in vitro* via the translocation of fluorescently labeled dextrans by adding them to the apical chamber of the transwell after 24 hours of incubation with the bacteria (*26*, *27*). We observed almost no translocation for any of the EcN derived strains, while the positive control (*S. typhimurium*) led to significant translocation of the polymer to the basolateral chamber (Figure 3C). While Caco-2 cells are a crude mimic of the mucosal epithelium, they are known to respond to exposure to pathogenic bacteria in predictable ways that include the activation of pro-inflammatory signaling cascades and release of cytokines such as IL-8 (*26, 28-31*). We monitored the response of polarized Caco-2 cells to apical bacterial exposure for 24 hours and found that EcN and the PBP8 variants showed no significant differences in IL-8 production, while *S. typhimurium* showed a 4-fold increase compared to cells with no bacterial exposure (Figure 3D). Overall, the transwell assays indicated that neither PBP8, nor the expression of wt-CsgA or CsgA-TFF3 led to any phenotypes that compromised epithelial integrity compared to unmodified EcN.

**Figure 3.**
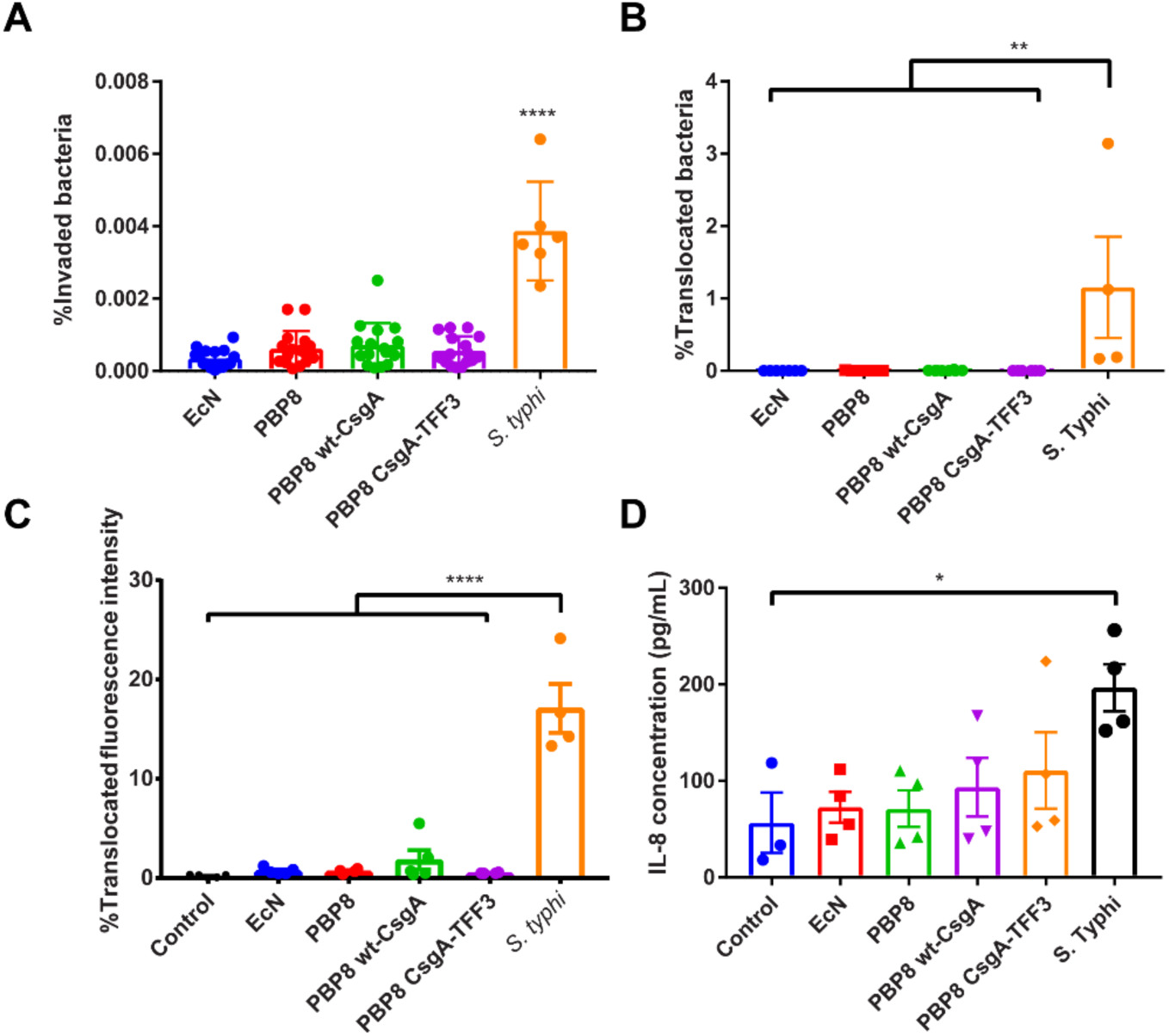
Effects of curli fiber expression on EcN pathogenicity. (A) Percent of bacteria that invaded a monolayer of Caco-2 after 2 hours of co-incubation with EcN, PBP8 variants, and S. typhimurium. (B) Bacterial translocation to the basolateral compartment of polarized Caco-2 cells exposed to bacterial library for 5 hours. (C) Epithelial permeability of polarized Caco-2 24 hours post-infection, quantified via FITC-dextran (MW 3,000-5000) translocation. (D) IL-8 secretion from the basolateral compartment of polarized Caco-2 cells 24 hours post-infection. Data are represented as mean ± SEM. See materials and methods for statistics.

### Engineered EcN can transiently colonize the mouse gut and express curli fibers in situ

Our initial *in vivo* experiments focused on demonstrating the viability and persistence of engineered EcN strains in the mouse GI tract after oral administration. For these experiments, we employed an EcN strain with a genomically integrated luminescence operon (*luxABCDE*) to facilitate *in vivo* tracking (*32*, *33*). After transformation of this EcN strain with the panel of plasmids encoding the synthetic curli operons, the strains were administered to healthy mice (C57BL/6NCrl) via oral gavage, concurrent with drinking water containing kanamycin and L-(+)-arabinose in order to maintain the plasmids and induce curli expression. A single dose of 10^8^ colony forming units (CFU) led to persistent colonization (>32 days) of the mouse GI tracts for all but one of the curli producing strains (EcN wt-CsgA, EcN CsgA-TFF2, and EcN CsgA-TFF3), as measured by CFU counted from fecal samples. Notably, EcN wt-CsgA maintained a CFU count of 10^8^-10^9^ over the course of the experiment, similar to that of EcN transformed with a pBbB8k plasmid encoding for green fluorescent protein (GFP) as a control (Figure 4A). This suggested that wild-type curli fiber overproduction *in vivo* did not compromise the fitness of the engineered EcN any more than recombinant production of any heterologous intracellular protein. In comparison, EcN CsgA-TFF2 and EcN CsgA-TFF3 concentrations fell over the course of the experiment to ~10^6^-10^7^ CFU, suggesting that production of the CsgA-TFF fusions was stressful enough to compromise the colonization ability of EcN in the stringent environment of the mouse gut. We speculate that this may be due to the large size of the TFF fusion domains (*34*), in addition to their propensity to form internal disulfide bonds that could hinder extracellular export. Indeed, the difference in growth rates between wt-CsgA and CsgA-TFF producing strains is reflected in the *in vitro* growth rates (Figure S3). We further confirmed the presence of the engineered EcN strains in living animals by visualization of the bacterial luminescence on days 2, 6, and 10 after oral administration using an *in vivo* imaging system (IVIS). As expected, the luminescence of the EcN could be observed in the abdomen of all mice that had received bacteria (Figure 4B and S4).

**Figure 4.**
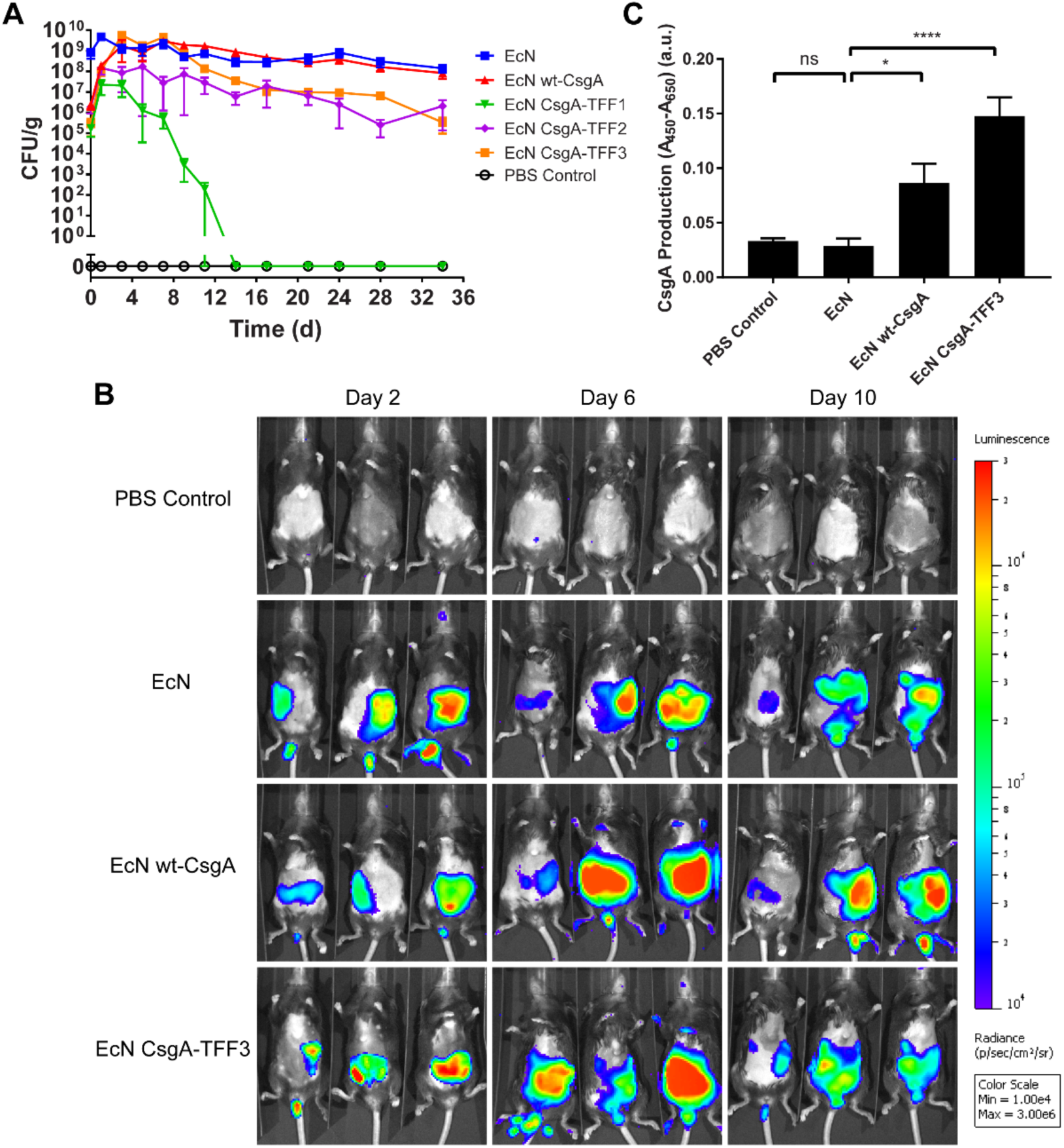
Residence time of engineered EcN in the mouse gut and in situ curli expression. (A) In vivo residence time for EcN variants, as measured by CFU counting from fecal samples. (B) IVIS images of mice that received either PBS or luminescent EcN variants at various time points post-inoculation. (C) Relative CsgA production from fecal sample analysis from mice administered 5 days post-inoculation, as measured by ELISA. Data are represented as mean ± SEM. See materials and methods for statistics.

In addition to the engineered bacteria themselves, we sought to confirm that the engineered curli fibers were being produced *in situ*. We therefore performed ELISA using an anti-6xHis antibody on homogenized fecal samples obtained 5 days after oral administration. We found that, as expected, only mice that had received curli producing strains (EcN wt-CsgA and EcN CsgA-TFF3) showed signal above background levels (Figure 4C). In comparison, EcN producing GFP only showed only background signal. Finally, we used immunohistochemistry at the experimental endpoint to visualize the engineered curli fibers directly in tissue sections of the proximal and distal colon (Figure 5). The curli fibers were probed with a fluorescently labeled anti-6xHis antibody (*magenta*), while anti-MUC2 (*red*), anti-E-cadherin (*green*), and Hoechst (*blue*) stains were used to visualize the mucins and colonic epithelial cells. We found that due to the fixing and staining protocol, which were designed to preserve the integrity of the intestinal mucins, the Hoechst and E-cadherin antibodies stained not only the colonic cells, but also DNA and cytoskeletal elements found in the feces and other contents of the gut lumen present in the tissue sections. Nevertheless, the distinct pattern and arrangement of colonic cells in the sections could be used to identify the epithelial border and differentiate them from luminal content. The anti-6xHis staining revealed that wt-CsgA could be observed throughout the gut lumen and near the most superficial layers of mucus, while the PBS control showed minimal background staining. Interestingly, the CsgA-TFF3 signal co-localized with the MUC2 signal, suggesting that the mucin binding activity of the TFFs promoted mucus integration of the engineered curli fibers.

**Figure 5.**
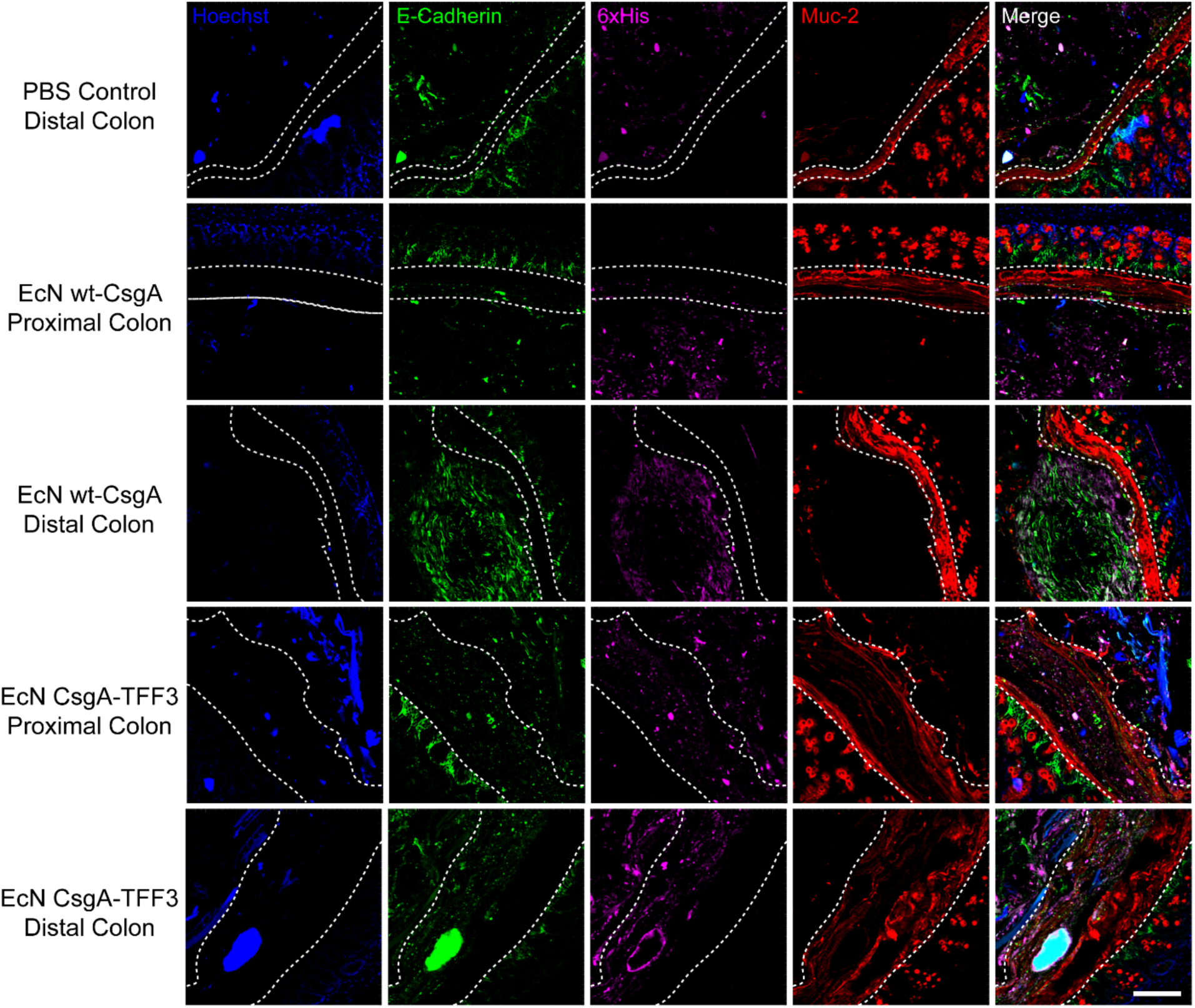
Immunohistological visualization of engineered curli fibers in tissue sections from proximal and distal colons of mice receiving different bacteria. Sectioning protocol was designed to preserve mucus and luminal content. Sections were stained with Hoescht (blue) and fluorescently labeled antibodies: anti-E-cadherin (green), anti-6xHis (magenta), anti-Muc2 (red). Last column shows an overlay of all stains. The white dotted lines represent the boundary of the epithelium and mucus layers (scale bar = 100 μm).

### Engineered EcN ameliorates disease activity in a mouse model of DSS-induced colitis

Based on the ability of the engineered EcN strains to colonize the mouse GI tract and the previously demonstrated wound healing and mucin binding activity of CsgA-TFF3 curli fibers *in vitro* (*22*), we sought to investigate their efficacy in a dextran sodium sulfate (DSS) model of murine colitis (*35*, *36*). DSS is a chemical colitogen that can be administered orally to induce epithelial damage and compromised barrier function in the mouse colon. Unmodified EcN and TFF secretion from *Lactococcus* spp. have each individually shown some efficacy against the DSS model, so we reasoned that their combination with the PATCH system would have similarly beneficial effects (*11*).

We examined the protective effects of engineered CsgA-TFF3 produced by PBP8 using female C57BL/6NCrl mice that had been administered 3% DSS over several days to induce colonic inflammation. Pilot experiments with oral administration of PBP8 strains were not very effective in decreasing disease symptoms. However, histological analysis and further literature consultation revealed that DSS induced colitis was most severe in the distal colon, whereas the engineered bacteria resided mostly in the cecum and proximal colon (Figure S5) (*24*, *37*, *38*). In order to circumvent this peculiarity of the murine DSS model and investigate the efficacy of our approach, we pivoted to rectal administration of the bacteria so that they could easily co-localize with the affected tissues. Notably, we do not envision that this issue would affect the efficacy of engineered bacteria in other models or in humans, as both oral and rectal deliveries are viable routes of drug administration depending on the patient’s disease localization.

As outlined in Figure 6A, the mice received daily administrations of PBP8 rectally for three days prior to DSS intake, during five days of DSS intake, and during a five day recovery period. DSS treatment in mice that had not received any bacteria (PBS DSS^+^) led to intestinal inflammation that could be observed by weight loss and increases in disease activity index (DAI, a composite measure of weight loss, diarrhea index and rectal bleeding, Table S1) compared to mice in a healthy control group without DSS treatment (PBS DSS^−^). Mice that received bacteria expressing CsgA-TFF3 (PBP8 CsgA-TFF3 DSS^+^) had significantly ameliorated weight loss and reduced DAI over the course of the experiment and almost returned to the same state as the healthy control group 5 days after DSS removal (Figure 6B,C). Mice that received engineered bacteria that were either producing cytosolic GFP as a control (PBP8 DSS^+^) or producing wild-type CsgA (PBP8 wt-CsgA DSS^+^) showed results similar to the disease group without any bacterial administration. The mice were sacrificed at day 10 for investigation of their colons to examine the effects of PBP8 CsgA-TFF3 on the induced colitis. DSS inflammation is associated with colon length reduction (*39*). We found that colon length did not differ significantly between the PBP8 CsgA-TFF3 group and the healthy control group, suggesting that *in situ* CsgA-TFF3 production attenuated colonic inflammation caused by DSS. In comparison, the colitic control, PBP8, and PBP8 wt-CsgA groups all had shorter colons (Figure 6D).

**Figure 6.**
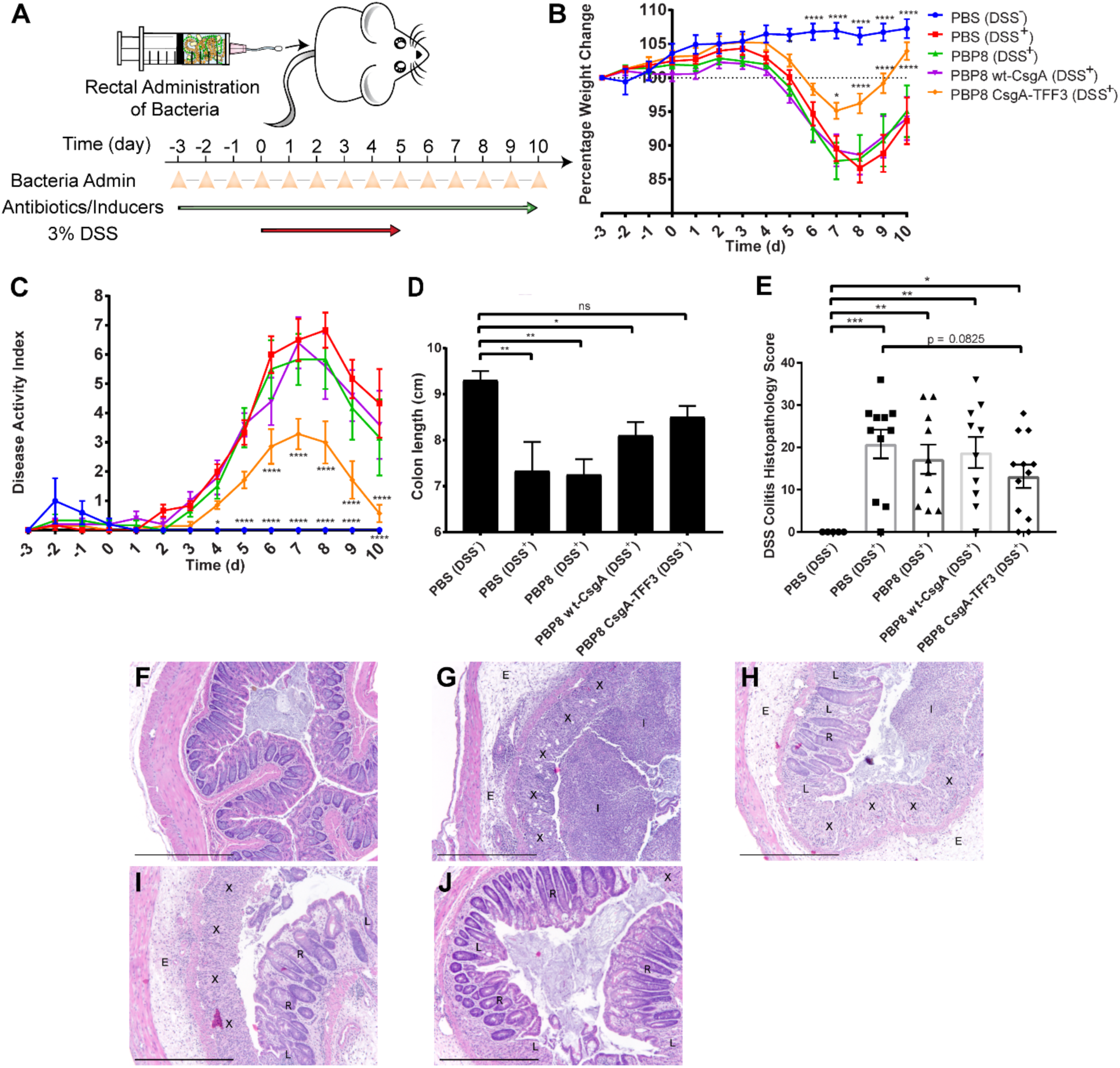
Therapeutic efficacy of engineered EcN against mouse model of DSS-induced colitis. (A) Schematic of administration schedule. PBP8 variants (10^8^ CFU) were administered rectally daily. Antibiotics and inducers were administered continuously via drinking water. Weight change (B, N = 9) and disease activity index (C, N = 5-7) over time, averaged across two independent experiments. Activity index criteria are described in Table S1. (D) Colon length at endpoint from two independent experiments (N = 5-7). (E) Combined DSS colitis histopathology score reflecting severity of inflammation (see Table S2 for details), at endpoint from two independent experiments (N = 5-10). (F-J) Representative histology of distal colon sections stained with haematoxylin and eosin from each experimental group: (F) PBS (DSS^−^) – healthy control, (G) PBS (DSS+) – colitic, (H) PBP8 (DSS+), (I) PBP8 wt-CsgA (DSS+), (J) PBP8 CsgA-TFF3 (DSS+). Image markers indicate complete loss of crypt and goblet cell depletion (X), immune cell infiltration (I), tissue edema (E), partial loss of the crypts (L), and recovery of crypts (R) (scale bar = 500 μm). Data are represented as mean ± SEM. See materials and methods for statistics.

We also assessed the effects of bacterial administration on gut inflammation using histology. Common histological characteristics of DSS induced colitis include immune cell infiltration involving multiple tissue layers as well as loss of colonic crypts and epithelial damage (*40*, *41*). We employed a histological scoring system based on previously published accounts (Table S2) (*40*-*42*). All of the groups that received DSS treatment had significantly higher histopathology scores than the healthy control group, indicating more inflammatory effects (Figure 6E). The histology score for the PBP8 CsgA-TFF3 group was the lowest of all the groups that received DSS, though the difference was not quite significant according to our scoring and statistics criteria. Images of the histology sections from the colitic control group showed complete loss of crypt structure, goblet cell depletion, immune cell infiltration into the lumen, and edema of the tissues around the colon, all of which reflect the severe inflammatory effects of DSS treatment (Figure 6F). In contrast, tissue sections from mice treated with bacteria showed some qualitative improvements, with better preservation of crypt structure. The PBP8 CsgA-TFF3 group also showed lower inflammatory cell infiltration, less edema, and more intact epithelium. One explanation for the histological similarity among the DSS treated groups could be that the tissue sections were obtained 5 days after DSS treatment had stopped. Therefore, natural healing processes could have obscured any quantitative differences in histopathology between the groups that received DSS and bacteria.

### Anti-inflammatory effects of engineered EcN are associated with immunomodulation and barrier function enhancement

In order to start probing the mechanism of the apparent protective effects of the PATCH system with CsgA-TFF3, we analyzed gene expression and protein production profiles from colonic tissue homogenates across the experimental groups. The DSS colitis model leads to several such changes, including downregulation of genes associated with epithelial barrier function and upregulation of genes associated with inflammatory signaling (*43*, *44*). With respect to tight junction protein-1 (TJP-1), a.k.a. zona occludens-1 (ZO-1), the PBP8 CsgA-TFF3 group showed significantly higher mRNA levels than the colitic control group (Figure 7A). This is in line with the known functions of TFFs in mammalian hosts even though other markers of barrier function associated with TFF bioactivity (occludin, claudin-2, and intestinal TFF3, Figure S6A-C) did not show changes according to qRT-PCR analysis. Notably, the PBP8 wt-CsgA group also exhibited high *tjp-1* mRNA levels. This could the explained by known interactions between wt-CsgA and toll-like receptor 2 (TLR2), which can indirectly lead to moderate increases in *tjp*-*1* expression (*45*, *46*). Although the effects of TFF3 fusion to CsgA on its interaction with host receptors is not clear form this work, it is possible that both domains could contribute to modulating local gene expression.

**Figure 7.**
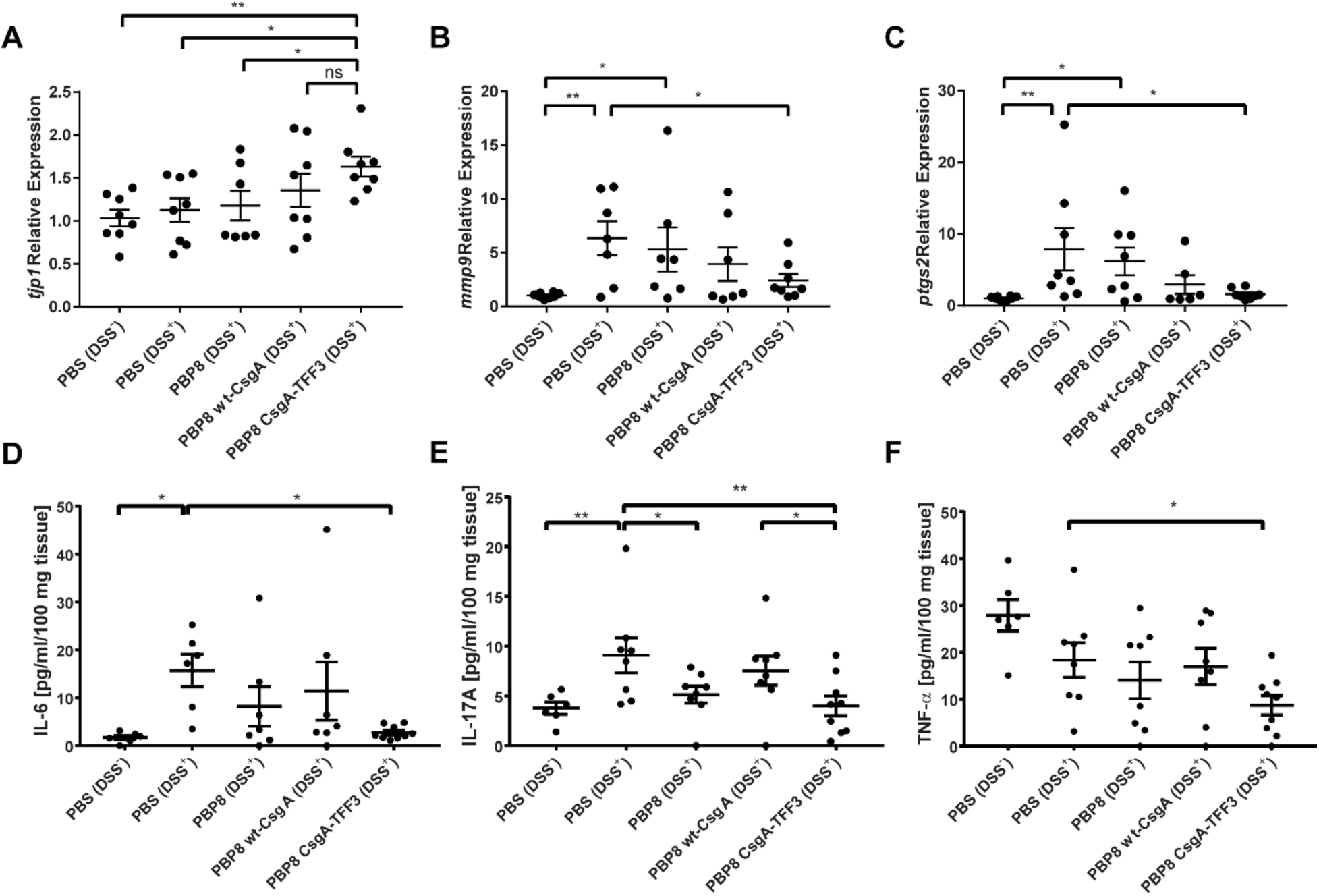
Effects of engineered EcN treatment on colonic gene and cytokine expression in DSS-induced colitic mice (A-C) tjp1, mmp9 and ptgs2 expression measured by qRT-PCR from homogenized distal colon sections. Data are presented as fold change compared to the healthy control group from two independent experiments (N = 7-9). (D-F) IL-6, IL-17A, and TNF-α protein level determined by multiplex ELISA. Data are presented as protein concentration per 100 mg of tissue from two independent experiments (N = 6-9). Data are represented as mean ± SEM. See materials and methods for statistics.

Matrix metalloproteinase-9 (MMP-9), whose function is to breakdown extracellular matrix proteins in inflammatory environments, is upregulated during IBD and is an essential mediator of tissue injury during DSS colitis. Blockade of MMP-9 has also been explored as a clinical treatment for IBD (*47*). In our experiments, the colitic control and PBP8 control groups showed elevated *mmp9* mRNA levels compared to the healthy control group, while the PBP8 CsgA-TFF3 group showed significantly lower *mmp9* expression (Figure 7B). Prostaglandin-endoperoxidase synthase 2 (PTGS-2), a.k.a. cyclooxygenase-2 (COX-2), showed similar results to MMP-9, with the PBP8 CsgA-TFF3 group significantly lower than the colitic group, and comparable with the healthy control group (Figure 7C). PTGS-2 is an enzyme whose production is induced in the colon during active IBD (*48*). Its inhibition has also been investigated as an anti-inflammatory treatment, and has ameliorated colitis in a murine DSS model (*49*). Similar to the case of MMP-9, the PBP8 wt-CsgA group also showed reduced *ptgs2* expression, which likely reflects the known interactions of unmodified curli fibers with the gut epithelium observed in this work and elsewhere (*50*).

Finally, we used a parallelized ELISA assay (Luminex), to probe cytokine levels from tissue homogenates obtained from mice at the experimental endpoint. We found that IL-6, IL-17A, and TNF-α concentrations decreased in colonic tissues for the PBP8 CsgA-TFF3 group compared to the colitic group (Figure 7D-F). IL-1B concentrations were also lower, but not statistically significant (Figure S6E). Cytokines such as TGF-β, IL-10 and IFN-γ were not significantly affected by any of the bacterial treatments (Figure S6D, F-G). The affected cytokines (IL-6, IL-17A, TNF-α and IL-1β) directly relate to differentiation and secretion of T helper 17 (Th17) cells, which is a lineage of CD4^+^ cells that mediates innate and adaptive immunity against various pathogens at mucosal sites (*51*).

## Discussion

We have developed an engineered beneficial bacterium that produces a self-assembled matrix *in situ* within the GI tract with programmable functions. Our engineering strategy (PATCH) enabled the production of a modified curli fiber matrix with fused TFFs under physiological conditions in a manner that does not appear to alter EcN’s inherent lack of pathogenicity. Administration of the engineered bacterium before, during, and after the induction of colitis in mice led to amelioration of inflammation and a reduction in Th17 responses in the colon. The protective effect of PATCH with CsgA-TFF3 also correlated with enhanced mucosal barrier function and is correlated with a reduction in the expression of inflammatory cytokines and enzymes in colonic tissues. Although more detailed studies will be required to rigorously confirm the mechanism of these effects, results from the genetic and biochemical analyses we performed were in line with previous accounts of trefoil factor bioactivity, which includes immunomodulation, the promotion of epithelial restitution, and upregulation of tight junction proteins (Figure 1) (*20*, *21*).

Ongoing work in the lab is focused on improving the PATCH system in order to make it more suitable for clinical deployment. In the experiments described here, antibiotics and small molecule inducers were fed to the mice in order to maintain the plasmid’s stability in the modified EcN strains and induce curli production, respectively. The antibiotics alter the gut microbiome significantly (*52*). To address this issue, second generation PATCH systems should be engineered with stable plasmid systems that do not require antibiotics that have been reported on extensively elsewhere (*33*, *53*, *54*). Although we demonstrated that oral administration was a viable approach to establishing a detectable and stable population of engineered EcN strains in the murine lower intestine, we ultimately opted for rectal administration to colocalize the bacteria near diseased areas and demonstrate efficacy. However, human IBD pathology can be spread across the colon and parts of the small intestine. Furthermore, E. coli is similarly localized in humans, and is known to proliferate at sites of inflammation (*55*). Therefore, either oral or rectal delivery may be relevant for other models of IBD or in humans. We also recognize that the curli fibers themselves are not a “blank slate” material in that CsgA already has numerous known interactions with host cells and tissues that could confound the effects of the displayed domains (*46*, *50*, *56*). While this may or may not impede further development of PATCH with curli fibers as a scaffold, we know that the biosynthetic machinery dedicated to curli secretion can tolerate a wide range of heterologous proteins (*57*). We are therefore in the process of exploring other combinations of scaffolding proteins and bioactive domains that can be secreted through the curli (a.k.a. “Type VIII”) pathway to circumvent these confounding effects and probe different therapeutic modalities.

Effective treatments for IBD with fewer undesirable side effects are a pressing medical need. Our platform has several features that make it potentially useful for clinical applications, including compatibility with oral administration, local biologic delivery that could mitigate systemic side effects, and high local concentrations of therapeutic entities in the gut lumen due to their anchoring to a self-assembled matrix. While further work is required to determine if this strategy can overcome the limited potency of other engineered microbial endeavors in human trials, it may work well as a combination therapy with other interventions for long-term disease management. Furthermore, the modularity of the PATCH system may make it useful for the display of domains with other therapeutic targets, including GI cancers, enzyme deficiencies, and pathogen sequestration.

## Materials and Methods

### Cell strains and plasmids

*E. coli* Nissle 1917 Prop-Luc strain (EcN *LuxABCDE*, erythromycin resistance) was kindly provided as a gift from Sangeeta Bhatia’s lab (Massachusetts Institute of Technology). *E. coli* PBP8 strain was derived from *E. coli* Nissle 1917 by genomic deletion of the curli operon (*24*). *S. typhimurium* (strain SL1344) was provided by Pam Silver’s lab (Harvard University).

The design and construction of the synthetic curli operon encoding plasmids are described in detail elsewhere (*24*). Briefly, a pBbB8k plasmid backbone contains the genes csgBA∗CEFG as a single cistron, controlled by the araBAD promoter, where A∗ indicates either wild-type or chimeric CsgA. Gene fragments encoding 6xHis tag modified TFF1-3 domains were cloned into these vectors to create pBbB8k-CsgA-TFF1, pBbB8k-CsgA-TFF2, and pBbB8k-CsgA-TFF3 (see supplementary document for CsgA fusion sequences and Figure S7 for pBbB8k-wt-CsgA plasmid map). The list of bacteria strains and plasmids can be found in Table S3. The list of reagents with distributers can be found in supplementary reagent list document. The details of *in vitro* curli expression can be found in additional materials and methods.

### Quantitative Congo Red binding assay

Based on established protocols (*22*, *24*, *58*), one mL of bacterial culture was pelleted and resuspended in a 0.025 mM a solution of Congo Red (CR) in phosphate buffered saline (PBS) for 10 minutes. After pelleting the cells again, the absorbance of the supernatant at 490 nm was measured using a microplate reader. Normalized curli fiber production was calculated by subtracting the measured absorbance value from that measured for 0.025 mM Congo Red in PBS and normalized by the OD_600_ of the original bacterial culture.

### Whole-cell filtration ELISA

Following an established protocol (*22*, *24*, *58*), the bacterial cultures were diluted to OD_600_ of 0.3 with tris-buffered saline (TBS). The specimens (200 μL) were transferred to a Multiscreen-GV 96-well filter plate, filtered, and washed with TBST buffer (TBS, 0.1% Tween-20). After blocking with with 1% bovine serum albumin (BSA) and 0.01% H2O2 in TBST for 1.5 hours at 37°C, and subsequent washing steps, 50 μL of anti-6xHis antibody-horseradish peroxidase (HRP) conjugate (1:200 dilution) was added to each well and incubated for 2 hours at 25°C. For the TFF3 antibody binding assay, samples were incubated with 50 μL of anti-TFF3 primary antibody (1:450 dilution) for 2 hours at 25°C and washed three times with TBST buffer, followed by incubation of 100 μL goat anti-mouse-HRP conjugated secondary antibody (1:5000 dilution) for 1 hour at 25°C and three subsequent washes. After the wash steps, Ultra-TMB (3,3’,5,5’-tetramethylbenzidine) ELISA substrate (100 μL) was added to each well and incubated for 10 minutes at 25°C. To stop the reaction, 50 μL of 2 M sulfuric acid was added to each well. 100 μL of the final reaction was transferred to 96 well plate and measured the absorbance at 450 and 650 nm. The relative amount of displayed peptide was measured by subtracting absorbance at 450 nm with absorbance 650 nm.

### Electron microscopy

200 μL of the testing cultures were filtered onto Nucleopore Track-Etched membranes (0.22 μm pore size) under vacuum and placed in fixative solution (2 % glutaraldehyde and 2 % paraformaldehyde in 0.1 M sodium cacodylate buffer) for 2 hours at room temperature. After fixation, the membranes were gently rinsed with water and subjected to an ethanol gradient (25, 50, 75, 100 and 100 % (v/v)) with a 15 minute incubation for each concentration. The samples were then transferred to a critical point dryer). The dried membranes were placed on SEM sample holders with carbon adhesives and sputter coated with 80:20 Pt/Pd (5 nm-thick). A Zeiss Ultra 55 Field Emission SEM was used to image the samples.

### Invasion assay

The assay was adapted from previously published protocol (*22*). In brief, Caco-2 cells at passage 5-15 were plated in 24-well plates at a density of 105 cells per well in 500 μL of regular cell culture media and grown to 90% confluency. The bacterial cultures were pelleted, washed with PBS and diluted to an OD_600_ of 0.5 in DMEM with 1 g/L glucose and 1% FBS. The Caco-2 cells were rinsed twice with PBS to remove the antibiotic before addition of the bacteria (500 μL). Bacteria were incubated for 2 hours with the Caco-2 cells and removed by aspiration. The Caco-2 cells were then washed twice with 500 μL PBS before receiving 500 μL of DMEM, 1 g/L glucose, 1% FBS and 100 μg/mL gentamicin. After 1 hour of incubation, the media were aspirated and replaced with 1 mL of 1% Triton-X. The cells were incubated with Triton-X at 37°C for 10 minutes and repeatedly pipetted to homogenize. Each well was serially diluted and plated on kanamycin plates to count the colony forming units (CFU) of bacteria that had invaded the Caco-2 cells.

### Translocation assay

This assay was adapted from a published protocol (*26*). EcN variants and *S. Typhimurium* SL 1344 (Silver lab, Harvard Medical School) were grown and induced, if applicable, in LB media with appropriate antibiotics, pelleted, washed with PBS and diluted to OD_600_ of 0.01 in DMEM with 1% glucose and 1% FBS. One day before the experiment, the culture media of polarized Caco-2 (See additional materials and methods) were switched to their corresponding non-antibiotic counterparts with 1% glucose and 1% FBS. 600 μL of diluted cultures were used to infect polarized Caco-2 apically. After 5 hours of incubation, 100 μL of media were collected from apical and basolateral sides of the transwell, serially diluted and plated on antibiotic selective plates to enumerate percentage of translocated bacteria.

### Epithelial integrity

The experiment was adapted from a published protocol (*26*). The epithelial integrity of polarized Caco-2 was determined by TEER values, and a fluorescein isothiocyanate-labeled dextran (FITC-dextran) translocation experiment. Prior to the infection, the TEER value of each polarized Caco-2 well was determined using Millicell ERS-2 Voltohmmeter. The cells were infected in the same manner as the translocation assay protocol. At 24 hours post infection, the TEER value was measured again to calculate the reduction of TEER. At the same time, 5 μL of 10 mg/mL FITC-dextran (average molecular weight 3-5 kDa) was added apically to each transwell. Two hours after the addition, 100 μL of media from the apical and basolateral sides were collected and transferred to a black, clear bottom, 96-well plate and the fluorescence intensity was measured using a plate reader at 485 nm excitation and 520 nm emission wavelength.

### IL-8 production

Polarized Caco-2 cells were infected in the same manner as the translocation assay. At 24 hours post infection, 100 μL of media from the basolateral side was collected and assayed to determine the IL-8 concentration using ELISA kit.

### In vivo residence time study

This protocol was approved by the Harvard Medical Area Standing Committee on Animals (HMA IACUC) (Ref. No. 05185). Female 8-to 9-week-old C57BL/6NCrl mice were randomly assigned into six groups that each received a different variant of EcN Prop-luc bacteria: GFP, wt-CsgA, CsgA-TFF1, CsgA-TFF2, CsgA-TFF3 and PBS control. Mice were subjected to 18 hours of fasting prior to the experiment, to remove the food in the upper GI tract, but received L-(+)-Arabinose (10 g/L) and kanamycin (1g/L) via drinking water. At the beginning of the experiment (day 0), the bacterial samples were prepared by culturing them to log phase and concentrating them to OD_600_ of 10 in 20% sucrose in PBS. Mice were given either 100 μL of PBS or bacterial suspensions based on their specified groups through oral gavage. Afterward, mice were returned to normal chow with L-(+)-Arabinose and kanamycin drinking water. At day 0 (5 hours after the administration), 1, 3, 5, 7, 9, 11, 14, 17, 21, 24, 28, and 34, fecal samples from each mouse were collected, weighed, serially diluted and plated on antibiotic selective plates (erythromycin and kanamycin) to enumerate resident CFU over time.

### In vivo imaging of engineered microbes

Mice were given an alfalfa-free chow for at least 5 days prior to the experiment to minimize autofluorescence. Mice were given the engineered EcN in the same manner as the residence time protocol in terms of special water and fasting regime. At day 2, 6, 10, mice were shaved at the abdominal area to help expose the GI tract to the IVIS machine. Mice were then imaged under anesthesia using IVIS Lumina II using luminescence filter, with field of view (FOV) = D (12.5 cm), fstop = 1 and large binning. Living Image software version 4.3.1/4.4 was used for image analysis. The detailed protocol for curli immunohistochemistry can be found in additional materials and methods.

### Fecal filtration ELISA

Homogenized fecal samples from day 5 of the residence time experiment were transferred onto the Multiscreen-GV 96-well filter plate (0.22 μm pore size). The volumes were normalized so that about 1.25 mg of fecal samples were on each membrane. Then, the filter plate underwent similar processes: blocking, incubating with anti-6xHis-HRP antibody, and interacting with the TMB substrate, as the whole cell ELISA protocol described above.

### Dextran Sodium Sulfate (DSS) model of mouse colitis and treatment protocol

Mice were randomly assigned to five groups: non-colitic (PBS DSS^−^), colitic (PBS DSS^+^), PBP8 with GFP control vector-treated (PBP8 DSS^+^), PBP8 with wild-type CsgA vector-treated (PBP8 wt-CsgA DSS^+^), and PBP8 with CsgA-TFF3 vector-treated (PBP8 CsgA-TFF3 DSS^+^) with n = 4-5 in each group for each set of experiments. During the course of one experiment (14 days), mice were fed with normal mouse chow *ad libitum* while animal body weight and water intake were evaluated daily. At the beginning of the experiment (day −3), mice in all groups started receiving 10 g/L L-(+)-Arabinose and 1 g/L kanamycin in drinking water. Meanwhile, the PBP8 bacteria cultures were grown, induced overnight, centrifuged and resuspended to an OD_600_ of 10 in 20% sucrose in PBS. Then, mice received 100 μL PBS (in PBS DSS-and PBS DSS^+^ groups), PBP8, PBP8 wt-CsgA and PBP8 CsgA-TFF3 by rectal administration once daily throughout the experimental period. Three days after the start of bacterial administration (day 0), colitis was induced by the addition of DSS (MW 40,000, Alfa Aesar) to a final concentration of 3% in the drinking water that also contained L-(+)-Arabinose and kanamycin. Mice in all groups except the non-colitic (PBS DSS-) group received DSS treatment for five days. After DSS removal (day 6), all mice were given the L-(+)-Arabinose and kanamycin water until day 10 when they were sacrificed.

The colon of each mouse was removed and its length measured. The feces were gently scraped off. The distal colon of each mouse was divided into three sections (about 1 cm each). The most proximal section was placed in RNAlater solution, frozen with liquid nitrogen and stored at −80°C until RNA extraction. The middle section was weighed, frozen with liquid nitrogen and stored at −80°C for protein quantification. The most distal section was fixed in 4% paraformaldehyde in PBS buffer overnight at 4°C for histological analysis. Detailed protocols for the determination of disease activity and histological studies can be found in additional materials and methods.

### Luminex multiplex immunoassay

500 μL of 1X mammalian cell lysis buffer and 5 mm-stainless steel beads were added to frozen tissue samples in 2 mL microcentrifuge tubes. The samples were then homogenized using TissueLyser LT at 50 Hz for 10 minutes at 4°C. Following homogenization, the samples were centrifuged at 10,000 g for 10 minutes at 4°C and the supernatants were transferred to new sample tubes. The samples were tested for six cytokines: IFN-γ, IL-1β, IL-6, IL-10, IL-17A, and TNF-α, using a Bio-Plex Pro Mouse Cytokine Th17 Panel A 6-Plex kit in accordance with the manufacturer’s protocol. The final 96-well plate was processed using the BioPlex 3D system and the concentration of each cytokine was determined using Bio-Plex Manager software.

### Gene expression analysis by qRT-PCR

RNAlater-stabilized tissues were subjected to total RNA extraction using the TissueLyser LT and RNeasy plus mini kit in accordance with the manufacturer’s protocol. RNA samples were eluted with 100 μL RNase free water provided from the kit. Following the elution, the concentration of RNA was determined by spectroscopy using the Nanodrop 2000c. 10 ng of RNA was analyzed using specific primers for each gene of interest (Table S4) and a KAPA SYBR FAST One-Step qRT-PCR kit with CFX96 real time PCR detection system in accordance with the manufacturer’s protocol. The Pfaffl method was used to normalize the expression result (*59*). In brief, the E-ΔΔCt value, where E was the primer efficiency of each primer pair and -ΔΔCt was difference in the average Ct value of the control groups and the sample, was used to transform the Ct values of each sample into the expression values. Then, the expression values of the housekeeping gene, glyceraldehyde-3-phosphate dehydrogenase (GAPDH), were used to normalize the expression values of each gene in this study.

### Statistics

The investigators were not blinded to the experimental conditions during experiments and outcome assessment, except for the pathohistology analysis. Data are presented as the arithmetic means plus or minus SEM. The data were analyzed using GraphPad Prism 7. The statistical significance of the bar and scatter bar plots were determined using one-way ANOVA followed by Dunnett’s (Figure 2, 4), Tukey’s (Figure 3) or Fischer LSD multiple comparison (Figure 6D, E, Figure 7). The time course experiments, such as percentage weight change and DAI, were analyzed using two-way ANOVA following by Dunnett’s multiple comparison. An associated probability (*p* value) of less than 0.05 was considered significant and given one star, whereas the *p* < 0.01 was given two stars, *p* < 0.001 was given three stars, *p* < 0.0001 was given four stars, and *p* > 0.05 was given “ns” accordingly.

### Data availability

The authors declare that all relevant data supporting the findings of this study are available within the article and its Supplementary Information Files or from the corresponding author on request.

## Acknowledgement

This work made use of the Harvard Digestive Diseases Center (HDDC), the Harvard Center for Nanoscale Systems (CNS) and Harvard Medical School ICCB-Longwood Screening Facility, and the Wyss Institute for Biologically Inspired Engineering. We would like to thank Trevor R. Nash, Frederick R. Ward, Amanda Graveline, Andyna Vernet, Frank Urena, Jessica J. Kim, Daniel Um, Mofeyifoluwa Edun, Frederic Vigneault, Elaine Lim, Thomas Ferrante, Garry Cuneo, Magdalena Kasendra, and Rachelle Prantil-Baun for their help. P.P. thankfully acknowledges the royal Thai government scholarship. D.B.C. gratefully acknowledges the National Institutes of Health grant (2T32CA009216-36). This work was supported by National Institutes of Health (1R01DK110770-01A1), the Blavatnik Biomedical Accelerator fund, and the Wyss Institute for Biologically Inspired Engineering.

## Author Contributions

P.P., A.D.-T., N.S.J. conceived the idea and designed the experiments. P.P. cloned CsgA variant plasmids. P.P. performed the curli characterization experiments and data analysis. P.P. and F.B. performed the *in vitro* tissue culture pathogenicity experiments and data analysis. P.P., A.D.-T., I.G. performed the DSS colitis mouse experiments and sample collections, and data analysis. P.P. performed the qRT-PCR and Luminex, and data analysis. D.B.C. performed the blinded histopathology scoring and P.P. performed the data analysis. P.P., A.D.-T. and I.G. performed the residence time experiment. P.P. and A.D.-T. performed the IVIS experiment. P.P. performed the fecal filtration ELISA and curli immunohistochemical experiment. The manuscript was written through contributions of all authors. The final version of the manuscript has been approved by all the authors.

